# Cryo-EM Structure of AAV2 Rep68 bound to integration site AAVS1: Insights into the mechanism of DNA melting

**DOI:** 10.1101/2024.04.02.587759

**Authors:** R. Jaiswal, V. Santosh, B Braud, A. Washington, Carlos R. Escalante

**Author notes:** Corresponding author **CRE Email:**.

## Abstract

The Rep68 protein from Adeno-Associated Virus (AAV) is a multifunctional SF3 helicase that performs most of the DNA transactions required for the viral life cycle. During AAV DNA replication, Rep68 assembles at the origin and catalyzes the DNA melting and nicking reactions during the hairpin rolling replication process to complete the second-strand synthesis of the AAV genome. Here, we report the Cryo-EM structures of Rep68 bound to double-stranded DNA (dsDNA) containing the sequence of the AAVS1 integration site in different nucleotide-bound states. In the apo state, Rep68 forms a heptameric complex around DNA, with three Origin Binding Domains (OBDs) bound to the Rep Binding Site (RBS) sequence and three other OBDs forming transient dimers with them. The AAA^+^ domains form an open ring with no interactions between subunits and with DNA. We hypothesize the heptameric quaternary structure is necessary to load onto dsDNA. In the ATPγS-bound state, a subset of three subunits binds the nucleotide, undergoing a large conformational change, inducing the formation of intersubunit interactions interaction and interaction with three consecutive DNA phosphate groups. Moreover, the induced conformational change positions three phenylalanine residues to come in close contact with the DNA backbone, producing a distortion in the DNA. We propose that the phenylalanine residues can potentially act as a hydrophobic wedge in the DNA melting process.

## Introduction

Parvoviruses are a large family of non-enveloped viruses with a single-stranded (ssDNA) genome typically between 4 to 6 kilobases long [1]. Their genomes contain palindromic sequences known as inverted terminal repeats (ITRs) on both ends that fold into double-stranded hairpin telomeres [1–5]. Among parvoviruses, the Adeno-Associated Virus (AAV) has become a model system for studying all aspects of parvoviral biology mechanisms due to its prominent role as a gene therapy vector and its ability to integrate site-specifically into human chromosome 19 [6–8]. As a dependovirus, AAV requires a helper virus such as adenovirus or herpes virus to initiate second strand DNA synthesis from the 3’ telomere [9, 10]. Without a helper virus such as adenovirus or herpes virus, AAV can establish a latent infection by integrating its genome into a particular locus on the q arm of chromosome 19 [13–17]. This DNA site, known as adeno-associated virus integration site 1 (AAVS1), contains the Rep Binding Site (RBS) and terminal resolution site (trs) elements that mirror the AAV ITR. [11–14]. With only two open reading frames (ORF), AAV produces nine transcripts generated using multiple promoters, alternative splicing, and various translation start sites. One of the ORFs generates four transcripts that produce four non-structural Rep proteins; the two large Reps (Rep78 and Rep68) are involved in every facet of the viral life cycle, including replication, while the small Reps (Rep52 and Rep40) work as molecular motors to package the empty capsid [15]. The large Rep proteins are critical players during the hairpin-rolling replication of the AAV genome. The current replication model proposes that after the host machinery uses the 3’-ITR as a primer to generate an intermediate duplex molecule with a closed-end (Figure S1)[2, 3, 16–18]. Replicating the 5’ telomere involves generating a new 3’ end through a site and strand-specific nicking at the trs site [19–22]. This process is initiated by Rep68/ Rep78 binding to the RBS sequence, melting double-stranded DNA, and catalyzing the endonuclease reaction on the ssDNA trs site [23].

Rep68/Rep78’s ability to catalyze such diverse reactions is through the cooperation of an N-terminal origin-binding and nuclease domain (OBD) from the HUH family and a unique SF3 helicase domain (HD) [24–30]. Previous studies have shown that AAV Rep proteins can form multiple quaternary states in solution, including octameric and heptameric rings. This property is distinct from other members of the SF3 family, which only form hexameric rings [31, 32]. Additionally, the Rep68 protein is influenced by its interaction with various DNA substrates [26–28]. To form a stable complex with RBS DNA sites, Rep68 needs the collaborative interaction of OBD, linker, and HD domains [27]. Previously, we found that Rep68 forms a heptameric complex with an AAVS1 DNA site using analytical centrifugation and negative staining transmission electron microscopy [33]. To characterize this complex’s structure and obtain insights into the assembly and DNA melting mechanism, we determined the Cryo-EM structures of the Rep68-AAVS1 complex in different nucleotide-bound states. The structures show that in the apo state, Rep68 forms a heptameric complex where only three OBD domains interact with the RBS DNA site. The remaining OBDs form transient dimers with the DNA bound OBDs. The HDs form a double-tier ring where the oligomerization domains (ODs) form a closed ring structure around DNA. The AAA^+^ domains barely make any interaction with DNA and do not interact with each other, but upon ATPγS binding, the AAA^+^ domains undergo a large conformational change, inducing both interaction with the DNA backbone and interaction with each other. Only three subunits bind ATPγS, while the other four remain in an apo-like state. The binding of ATPγS also positions three phenylalanine residues (F364) to come in close contact with the DNA backbone, producing a DNA distortion. Mutation of F364 to alanine has a drastic effect on both DNA unwinding and trs nicking. We hypothesize that these residues may serve as a hydrophobic wedge to initiate DNA melting.

## Results and Discussion

### Overall Structure of the apo Rep68-AAVS1 complex

We used a truncated form of Rep68 (1-490), and a 50-bb long dsDNA substrate derived from the AAVS1 sequence (Figure 1A). The micrographs and 2D class averages clearly show structural features for a heptameric ring surrounding a central density representing DNA (Figure 1B-C). We determined the structure of the Rep68-AAVS1 complex in the absence of nucleotides following the data analysis workflow illustrated in supplementary Figure S2 to a final resolution of 5.3 Å. The reconstructed map, although low resolution, is of enough quality to show the overall architecture of the complex unambiguously. Seven helicase domains (A-G) form a discontinuous ring, while the density in the central channel of the ring corresponds to the AAVS1 DNA molecule, which spans approximately 40 base pairs (Figure 1D-F). Six OBD densities can be identified, of which three are DNA-associated. The remaining three form dimers with the bound OBDs, although these densities are less defined. The heptameric HD ring consists of two tiers representing the two domains of the SF3 helicase domain (Figure 1E). The OD tier density closely encircles the DNA, showing four helices forming the OD helical bundle in each of the seven subunits (Figure 3A). The density of the AAA^+^ domain ring is less defined and only shows the overall domain density, suggesting high mobility. The map shows that the DNA molecule is tilted toward one side of the AAA^+^ ring containing subunits E, F, and G.

**Figure 1.**
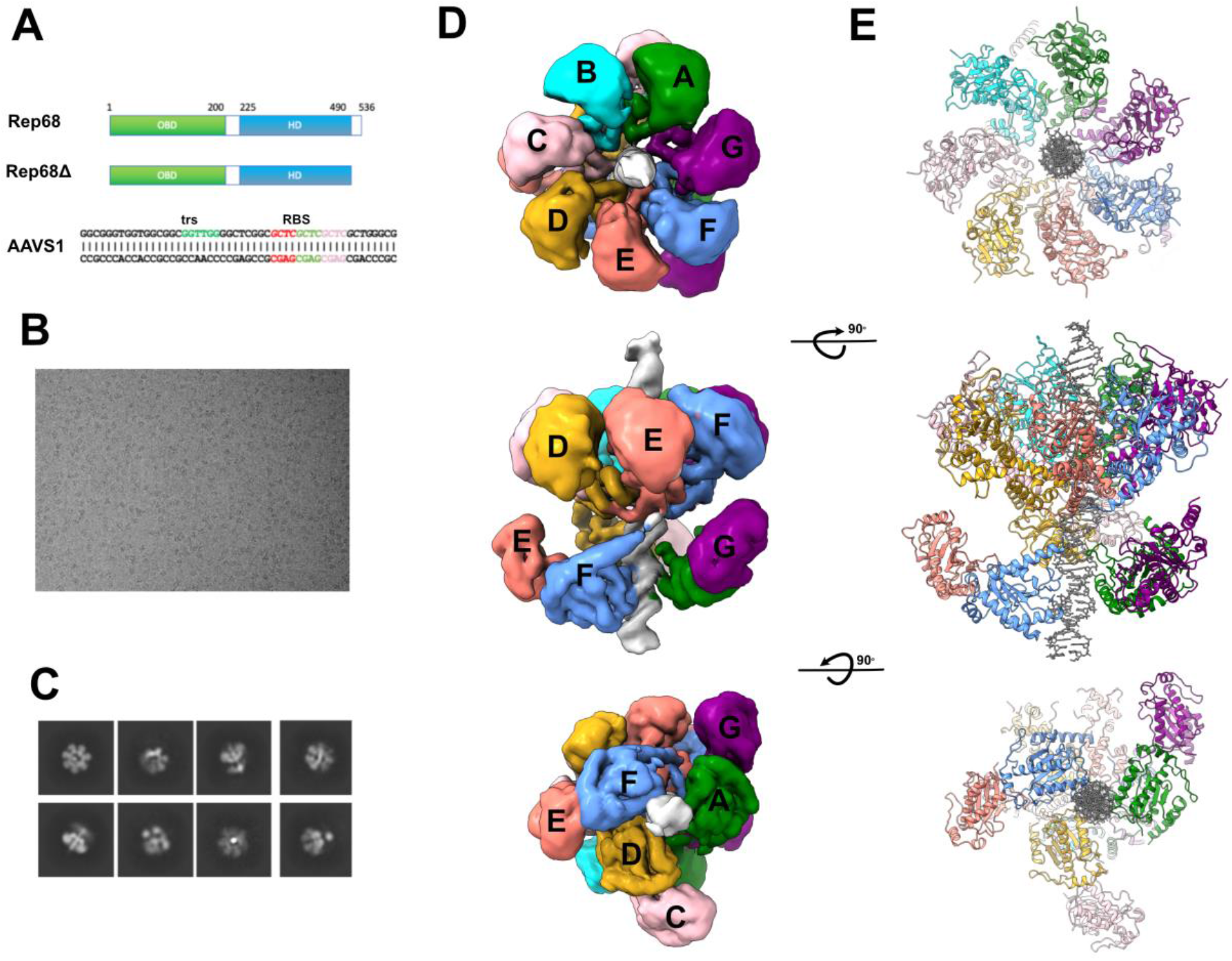
Cryo-EM Structure of Rep68-AAVS1. (A) Top, The schematic representation of the Rep68 construct shows the origin binding domain (OBD) in green and the helicase domain (HD) in blue. Middle, Construct used for these studies. Bottom, shows the sequence of the AAVS1 DNA site with the Rep Binding Site (RBS) and terminal resolution site (rbs) in red. (B) representative cryo-EM micrograph of Rep68-AAVS1 complex. (C) Typical 2D classes. (D) Top, Cryo-EM map viewed down the DNA axis with the HDs on front. Sideview of the overall map and bottom, view with OBDs on front. The subunits are colored with A (green), B (cyan), C (pink), D (goldenrod), E (salmon), F (cornflower blue) and G (purple). DNA is colored light grey. (E) Ribbon representation of the Rep68-AAVS1 complex viewed as in (D).

### Overall Structure of ATPγS bound Rep68-AAVS1 complex

We determined the structure of the Rep68-AAVS1 complex in the presence of ATPγS to a final resolution of 3.4 Å (Figure 2, S3). Similarly to the apo complex, the map shows an HD heptameric ring with dsDNA density in the middle of the ring spanning ∼ 28 bp (Figure 2A, B). We could only identify partial densities corresponding to one or two OBDs, but they are not well-defined due to their high mobility (Supplementary Figure S3). The heptameric ring is asymmetric and can be divided into two halves according to the interaction between subunits, and local resolution; the first hemi-ring, with higher resolution, is composed of three HDs (E, F, G) forming a continuous surface through both their AAA^+^ and OD domains (Figure 2A). The second hemi-ring is discontinuous, containing four HDs (A, B, C, D) with the AAA^+^ domains separated from the neighboring subunits, except for D, which loosely interacts with subunit E. (Figure 2A-C). All three subunits in the first hemi-ring have an ATPγS molecule bound and make subunit interactions with their neighboring subunits (Figure 2B). The consequence of ATPγS binding to only three of the subunits in the complex is striking in terms of the local resolution of the two hemi-rings, as shown in Figure 2C. This effect combines the stabilization of the AAA^+^ domain by the newly formed inter-subunit interactions and the more intimate binding to DNA. After sharpening the map with DeepEMhancer, we can identify two OBDs still bound to DNA. However, the density is too undefined to dock any models, alluding to their mobility [34].

**Figure 2.**
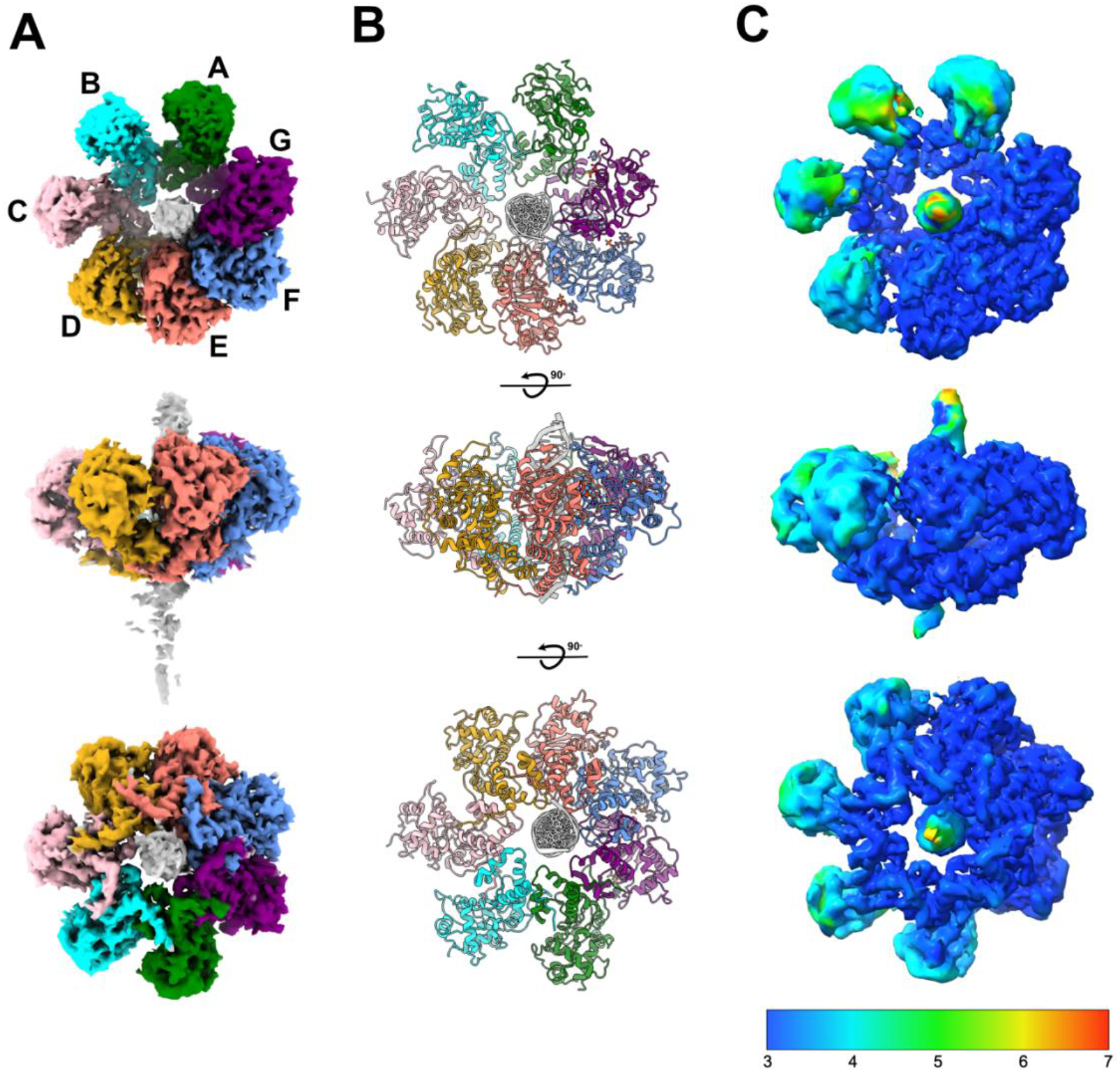
Cryo-EM structure of Rep68-AAVS1-ATPγS complex. (A) Top (HDs), side and bottom (OBDs) views of the Cryo-EM density. (B) Ribbon representation of the refined model. (C) Final map colored according to local resolution calculated in cryoSPARC.

### Interfaces between Rep68 subunits

#### Apo-complex

The complex is primarily stabilized by two interactions: the binding of three OBDs to DNA and the ODs forming a closed ring around it (Figure 1F, 3A). The formation of dimers among OBD domains was unexpected. Variability analysis indicates that this interaction is dynamic, as seen by the less-than-optimal density in these domains. Interestingly, the dimeric OBD-OBD interface is similar to that in the double-octameric Rep68-ssDNA complex [35]. Namely, α-helices B, and C, and the loop connecting strands β2 and β3 from the DNA-bound OBD interacts with α-helix D and loop L_DB_ of the second OBD molecule. The OBD dimers occur between subunits A/G, C/D, and E/F, with the density of the OBD of subunit B missing, indicating its high mobility (Figure 3A). Surprisingly, the contribution of the AAA^+^ domains to oligomerization is marginal; Figure 3B shows that the buried surface area (BSA) between the AAA^+^ domains is negligible; in contrast, each OD interface contributes approximately 750 Å².

**Figure 3.**
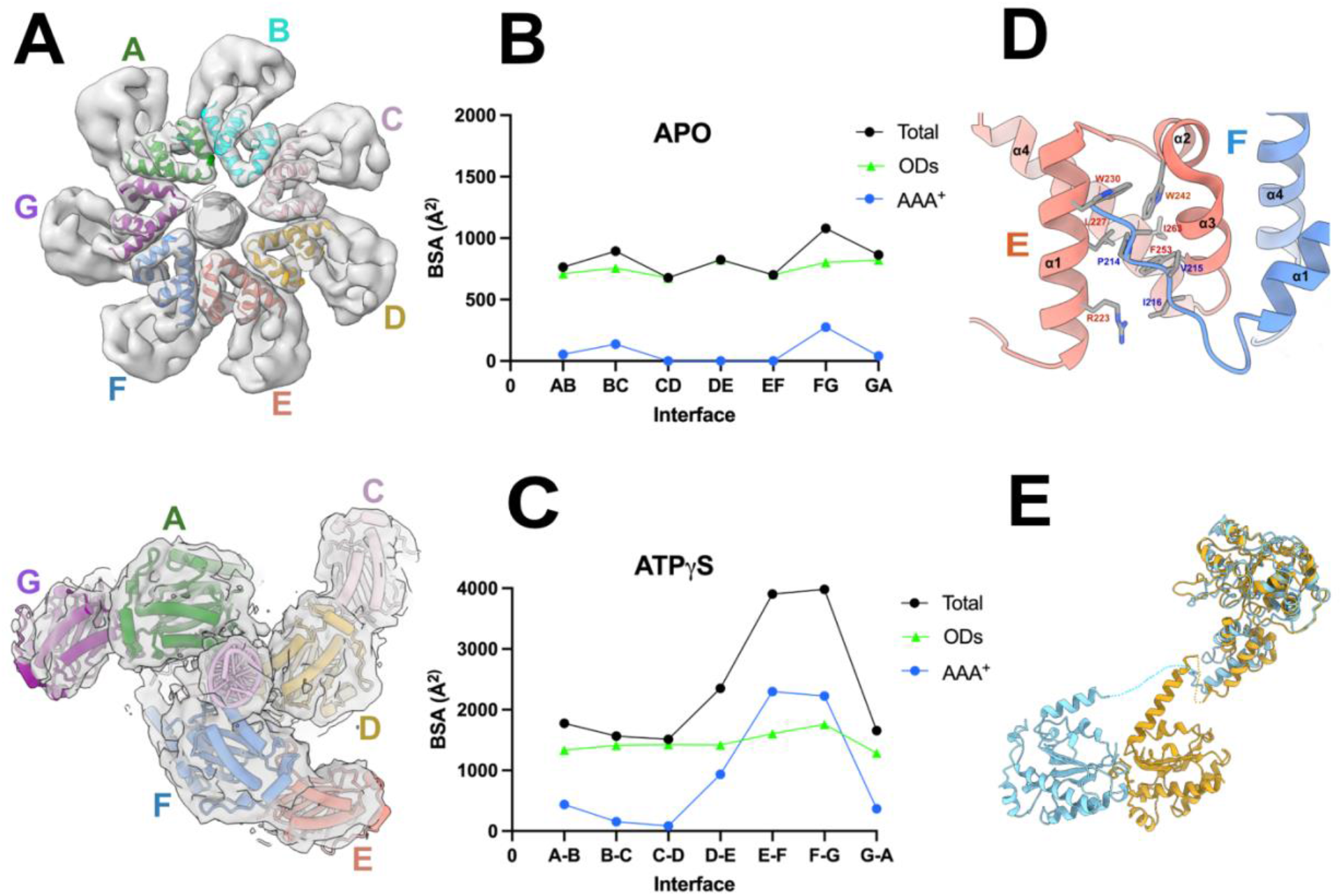
Interactions between Rep68 subunits. (A) Cryo-EM density shows the OD collar subunit arrangements with the docked ribbon representation. Bottom, OBD-OBD dimer interactions docket into the cryo-EM density map. (B) Plot showing the buried surface area (BSA) in each interface between subunits in heptameric complex. The BSA contribution of each of the HD subdomains OD (green triangles), AAA^+^ (blue), total (black circles) in the apo complex. (D) Plot showing the buried surface area of each of the subunit interfaces in the helicase domain (HD) and the contribution of each of the HD subdomains OD (green triangles), AAA^+^ (blue), total (black circles) in the ATPγS complex. (D) Representation of the linker ‘latch’ interacting with helices α1, α3 and α4 of the neighboring OD domain. (E) Superposition of subunits D and F illustrates the different linker distances between the OBD C-terminal helix and helix α1 of the HD.

#### ATPγS Structure

The binding of ATPγS leads to a nearly 1000 Å^2^ increase in the contribution of AAA^+^ domains to the heptameric ring’s total buried surface area. At the same time, there is an increase in the BSA of the ODs by about ∼200 Å^2^ compared to the apo complex (Figure 3C). We previously showed in the Rep68-ssDNA structure that the main contributor to the stability of the heptameric ring is a portion of the linker region that latches onto the neighboring subunit [35]. The higher resolution of the ATPγS complex Cryo-EM map enabled building the linker region from residues 213 to 224 (Figure 3D). The first half of this region (213-218) forms the ‘hydrophobic latch’ that engages in inter-subunit interactions with the OD of the neighboring subunit, as we previously showed [35]. Residues A213, P214, I215, and V216 stack against a crevice formed by α-helices 1-3 of the neighboring OD (residues R223, L227, W230, W242, F253, I263) (Figure 3D). Residue P214 plays an important role by changing the direction of the polypeptide chain and preventing clashes with the neighboring subunit. Residues 219 to 224 form an α-helix extending into the first helix of the HD (Figure 3D). Due to their inherent dynamic behavior, the linker regions preceding residue 212 connecting with the OBD domains are not visible on the map. However, this region needs to adopt numerous diverse conformations to accommodate the different distances between the OBD and HD domains around the complex, ranging from a compact one where the distance between the last and first residues in the domains is only 15 Å (subunit D) to highly extended ones in subunit E where the distance is 34 Å (Figure 3E).

### DNA induces large rigid body movements in the Helicase domain

Although the apo CryoEM map is not at high resolution, it is evident that even in the apo state, the AAA^+^ domains undergo varying degrees of motion around the ring, primarily in the form of a rigid body rotation of the AAA^+^ domain (Figure 4A). To determine the extent of the rotation, we aligned all subunits with the apo Rep40 X-ray structure (PDB id 1u0j) using the ODs as a reference [26]. In the apo complex, the rotation across the different subunits ranges from 15 to 51 degrees (Figure 4B). Interestingly, the subunits located farther from the DNA exhibit smaller rigid body rotations, while those closer to the DNA undergo a larger rotation (Figure 4B-C). This correlation is shown by plotting the degree of the AAA^+^ domain rotation as a function of the Rep68-DNA interface buried surface area. The data presented in Figure 4D indicate that in each complex, the 7 subunits fall into two different groups: one group has the four subunits that are farther away from DNA and exhibiting the lowest rigid body rotation, while the second group has the three subunits that have closer contact with DNA, suffering the largest rotations. Thus, binding to DNA induces rigid body motions in the AAA^+^ domains even without ATP binding. To determine the effect upon ATPγS binding, we performed the same analysis and observed that the AAA^+^ domains in subunits E, F, and G undergo additional rotation in the direction of the DNA molecule by an additional 5-15° (Figure 4D). At the same time, subunits A-C located in the opposite DNA face are further away from the DNA, causing minimal rotation (Figure 4D). Aligning the AAA^+^ domains among subunits E, F, and G shows no major differences between the functional motifs (rmsd avg 0.6 Å^2^), particularly in the pre-sensor-1 β-Hairpin (PS1βH). The structures demonstrate that only after ATP binding, the conformational changes are sufficient to promote direct interactions with DNA and among AAA^+^ domains.

**Figure 4.**
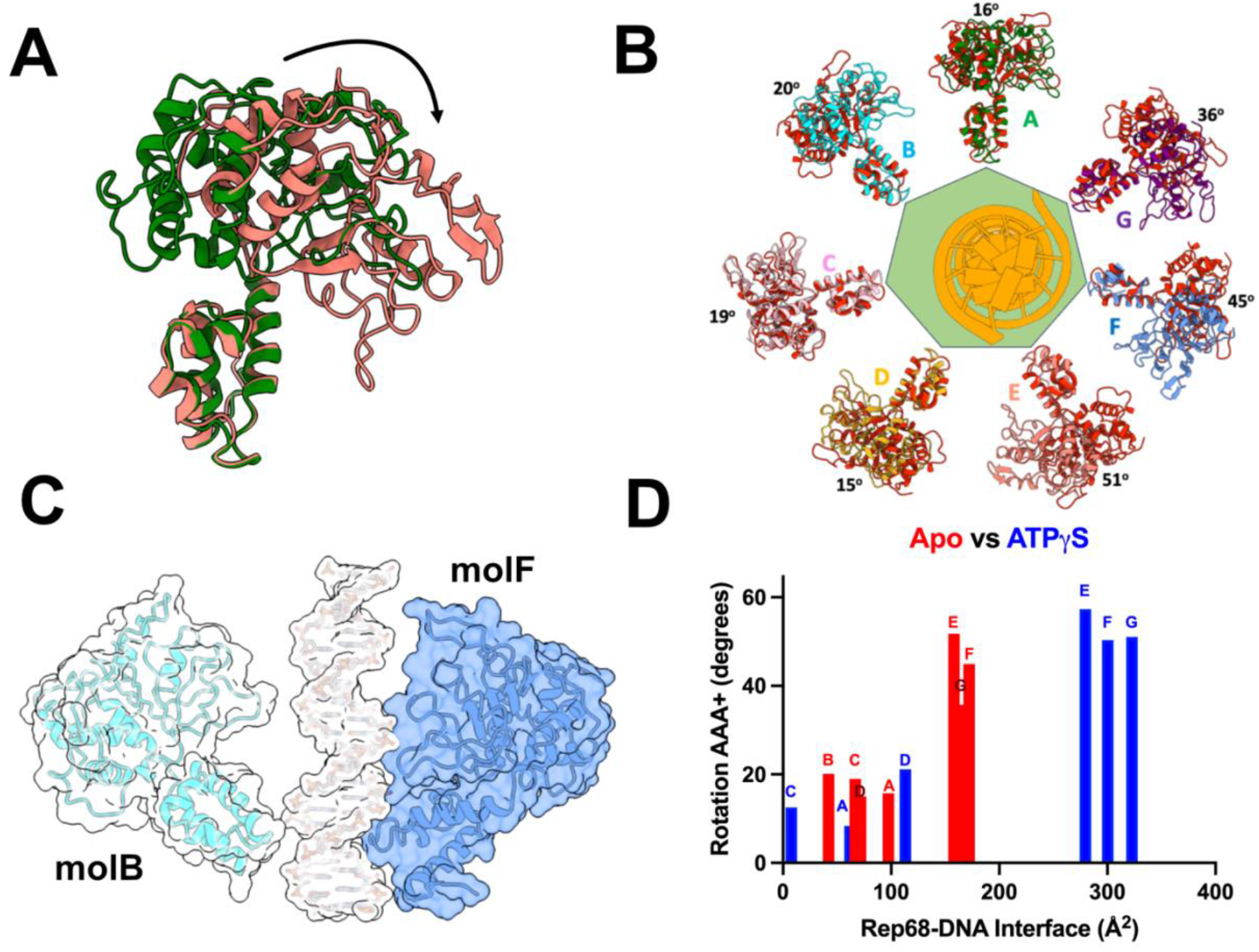
Conformation changes of the AAA^+^ subdomain. (A) Superposition of apo Rep68 HD subunits A and E. (B) Conformational changes around the DNA showing the rigid body rotation of the AAA^+^ subdomain when superimposed to apo Rep40. (C) Interactions of Rep68 subunits with DNA in opposite faces of the ring. (D) Plot of the AAA* rotation as a function of the Rep68-DNA interface buried surface area (BSA). The red bars correspond to the apo complex, blue bars are the ATPγS complex.

### Nucleotide states around the heptameric ring

We observe three different nucleotide states in the ATPγS complex; subunits A to D are in an empty (E) state with no nucleotide bound. Because the resolution in these subunits is between 6-7 Å, we based this assertion on the subunits’ overall characteristics, resembling those in the apo complex, particularly the small BSA between A-B, B-C and C-D interfaces (Figure 1, 3B, 3C). Subunit G is in a non-catalytic ATP-bound state (T) as subunit A is not close enough to form an interface (BSA ∼ 185 Å^2^), and the arginine finger (R444) is too far away to interact with the ATPγS molecule (Figure 5A, 5B). However, the nucleotide only interacts with the P-loop main chain atoms of residues T337, T338, and N342, explaining the higher B-factor in the nucleotide and the partial density in the base (Figure 5B). Only subunits E and F have all the required network of interactions to hydrolyze ATP and are in a catalytically competent T* state (Figure 5C, D). In addition to the P-loop residues, Walker A residues K340 and T341 interact with the nucleotide, and Walker B residues E378 and E379 are at a distance to coordinate with the Mg^+2^ ion. More importantly, the γ-phosphate interacts with the arginine finger residue R444. The conformation of each of the three ATPγS molecules has the adenine ring in a conformation similar to the ADP molecule seen in the X-ray structure of the Rep40-ADP complex (pdbid 1u0j) (Figure 5B-D) [36]. This conformation aligns the hydroxyl groups of the ribose ring to form hydrogen bonds with the main chain atoms of D455 in the nucleotide-binding loop (NB-loop) (Figure 5C, D). The interaction induces a conformational change in the NB loop that is crucial for promoting protein-protein interactions with the nearby subunit; particularly, it positions residue H482 of one subunit with H456 of the neighboring subunit to make π-π interactions (Figure 5C, D). Of note is that the D-E interface has the third largest BSA with ∼934 Å^2^ and subunit D could potentially have a nucleotide bound. However, the Cryo-EM map is not clear enough to confidently verify its presence and type of nucleotide even after processing the map with DeepEMhancer and local refinement [34]. Overall, the results show that ATP binding leads to interface formation among AAA+ subunits E, F, and G. However, only E and F subunits exhibit catalytic competence (Figure 5E).

**Figure 5.**
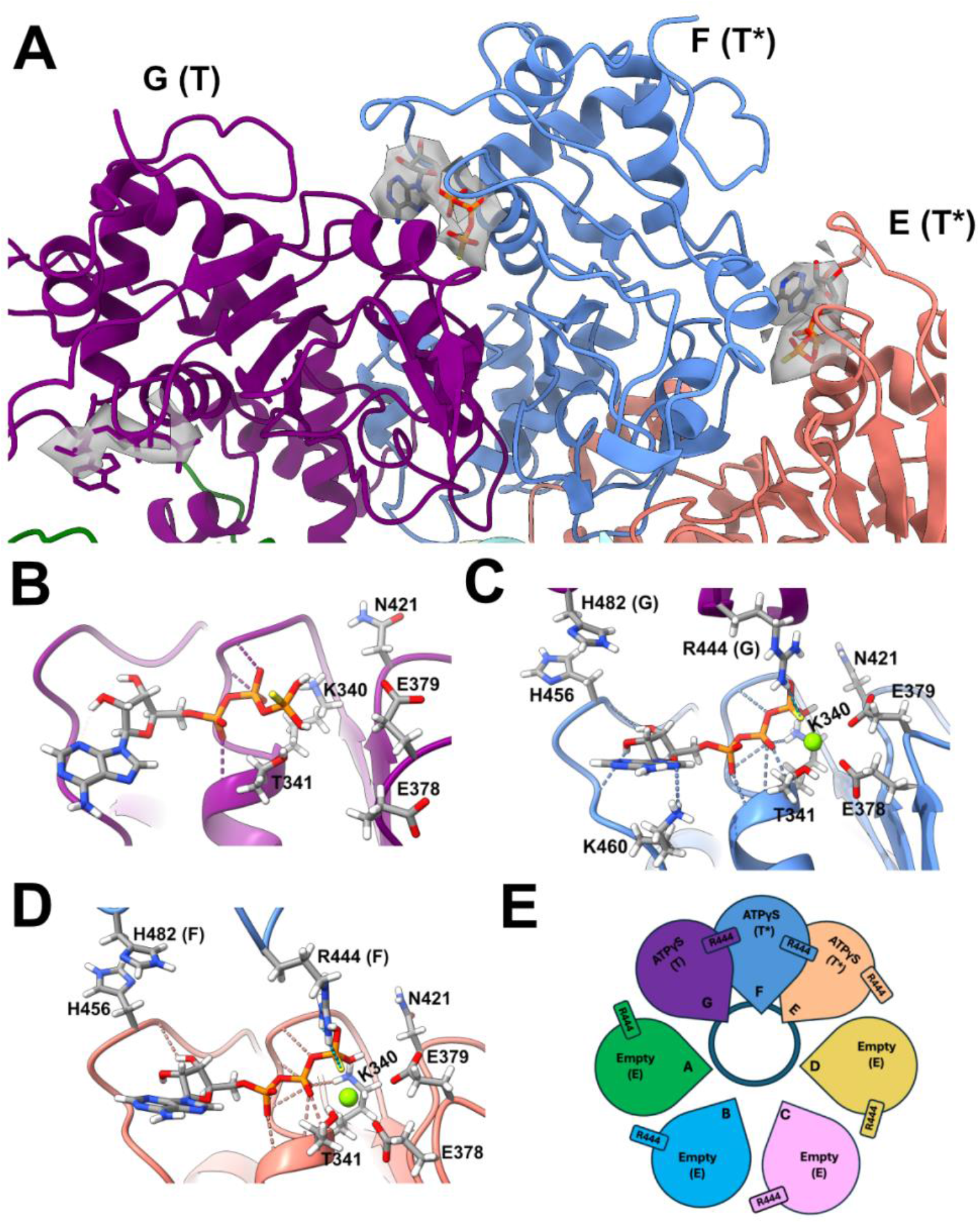
Nucleotide states around the heptameric ring. (A) Shown are subunits D (goldenrod), E (salmon), F (cornflower blue) and G (pink). The cryo-EM density of the nucleotide molecules is shown in transparent gray. (B) ATPγS binding pocket between subunits G-A, showing representative side chains interacting with the nucleotide. (C) ATPγS binding pocket between subunits F-G and (D) ATPγS binding pocket of E-F interface. Representative side chains are the arginine finger (R444), the Walker A residues K340 and T341 and the interface residues H456 and H482. (E) Nucleotide states around the Rep68 AAA+ ring showing that only in subunits F and G the argininine finger (R444) makes interaction to promote hydrolysis. The circle in the midle represent the DNA molecule.

### Rep68-DNA interaction

#### Apo Structure

The docking of the trimeric X-ray structure of the OBD-AAVS1 complex (PBD ID 4zq9) into the cryo-EM map of the apo complex allowed the determination of the register of the DNA sequence in the model (Figure 1D, 6A). Thus, in the apo complex the Cryo-EM DNA density covers the entire RBS-trs sequence (∼ 40 bp), showing the trs region is inside the top of the HD ring (Figure 6A). The structure shows that although the complex is heptameric, the maximum number of OBDs that can bind the RBS DNA region without steric clashes is three. Due to the insufficient resolution of the Cryo-EM map in the apo complex, the side chains involved in DNA interactions cannot be observed; however, because the docked X-ray structure of the OBD-AAVS1 complex was a perfect fit, we assume the recognition of DNA by the OBDs will be identical to those found in the X-ray structure [37]. The backbones of the seven OD domains are well defined in the map, showing they are close enough to interact with the DNA sugar-phosphate backbone via Van der Wall interactions using residues from the loop connecting helices α3 and α4, as this loop docks into the major groove (Figure 3A). The AAA^+^ domains do not appear to participate in direct interactions with the DNA; however, the AAVS1 DNA is tilted toward 3 of the subunits, and there may be some Van der Wall interactions, particularly in subunits F and G, via the PS1βH motif residues.

**Figure 6.**
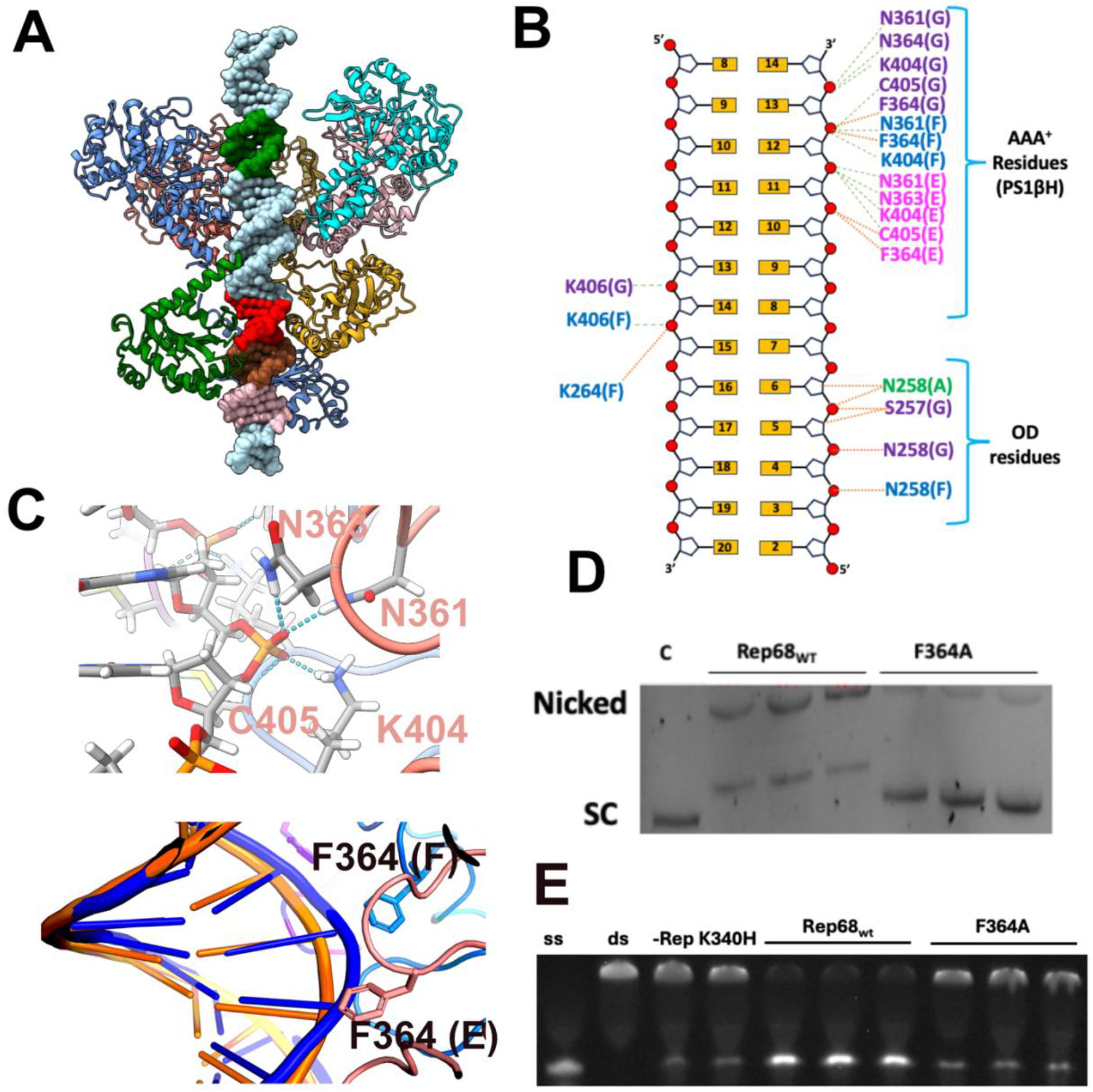
Rep68-AAVS1 interactions. (A) Overview of the Apo complex showing the span of the Rep68 ring and the position of the RBS and trs regions. For simplicity subunits B and C were removed. The RBS three GCTC repeats are colored red, brown and pink; trs in green. (B) Diagram showing the interactions of the Rep68 heptameric HD with DNA. (C) Representation of the interactions of the PS1βH and ϕ-loop with DNA of the E subunit. (D) Position of the three F364 residues along the DNA backbone shows the DNA distortion. (E) DNA gel showing results of plasmid nicking relaxation assay. C is the control supercoiled (SC) plasmid. Triplicate experiments of Rep68_wt_ and Rep68F364A. (F) Helicase activity assay lane 1, fluorescein-labeled-ssDNA, lane 2, fluorescein-labeled-duplex, lane 3, Rep68-K340H, lane 4-6, Rep68_wt_, lane 7-9, Rep68F364A.

#### ATPγS Structure

The ATPγS-bound state structure shows a paucity of DNA backbone interactions compared to other ss- or dsDNA translocases, where all subunits are engaged with DNA during the ATP hydrolysis cycle. Residues N361, N363, K404 and C405 from subunits E, F, and G make hydrogen bond contacts with three consecutive phosphate oxygens (Figure 6B). Most of these interactions occur in the same strand and are close to the trs region (Figure 6A). The interaction of C405 with the phosphate is through the main chain carbonyl in subunits F and G. Three consecutive F364 residues come into Van der Waals contact with the phosphodiester backbone, causing a small distortion in the DNA (Figure 6C). DNA parameter shows a widening of the major grove around this area compared to a B-shaped DNA molecule. A second area of distortion occurs around the OD tier (Figure S4). To study the impact of an F364A mutation on DNA melting, we employed the plasmid relaxation assay, which indirectly measures DNA melting necessary for trs nicking. Figure 6D shows that the nicking activity of the F364 mutant is severely impaired when compared to the WT protein, suggesting that F364 is necessary but not sufficient for DNA melting. After testing the helicase activity of the F364A mutant, we found that Rep68’s ability to unwind DNA is almost completely abolished by this mutation and has a severe impact on AAV-2 infective particle production (Figure 6D) [38].

## Discussion

This study shows for the first time how a full-length SF3 helicase assembles onto dsDNA and provides insights into the mechanism of how the large AAV Rep proteins may initiate DNA melting. The apo structure presented here shows that Rep68 forms a heptameric ring with dsDNA. However, only three OBDs remain bound to the RBS DNA site after assembly. Our results show that OD ring formation and the OBD-DNA interactions stabilize the complex. The heptameric ‘OD collar’ is formed by inter-subunit interactions of neighboring subunits using the 5-residue long hydrophobic latch [35]. The remaining linker residues connecting to the OBD display various conformations depending on the distance between the OBD and HD in each subunit, varying between 15 to as much as 34 Å (Figure 4D). This flexibility allows three OBDs to form transient dimers with the three OBDs bound to DNA (Figure 4A). A potential role for this interaction may be to prevent steric clashes among the DNA-unbound OBDs. Our results show that the OBD and OD almost work as a single unit to form the complex and bind DNA during assembly. This arrangement resembles the RepB plasmid initiator protein composed of only the OBD and OD domains [39].

The heptameric nature of the complex is unusual in SF3 helicases, which typically only form hexamers [27, 40, 41]; however, Rep68 can also form hexameric rings, but these have only been observed in the presence of ATPγS and ssDNA [35]. Analysis of the Rep68 hexamer inner channel dimension shows it is too narrow to fit dsDNA and that only the heptameric complex can fit a dsDNA molecule. We can look to other ring helicases that form mixtures of hexamers and heptamers to understand a potential role for this structure. In the bacteriophage T7 gene 4 Primase/Helicase, the human mitochondrial helicase Twinkle, and the HerA helicase from *Methanobacterium thermoautotrophicum*, the heptamer could enable loading onto dsDNA transition into an active replicative hexameric helicase [42–45]. We hypothesize that the Rep68 heptamer may play a similar role; whether it converts into a hexameric ring after or during DNA melting is still an open question.

A remaining question is how the coupling of ATP hydrolysis and DNA binding observed in the Rep68-AAVS1 complex leads to dsDNA melting. Our results show that upon binding ATPγS, the AAA^+^ domains undergo a substantial rigid body motion that promotes two events: the interaction of the AAA^+^ domain with DNA and the formation of AAA^+^ intersubunit interactions. Two motifs participate in DNA interactions: the SF3 signature PS1βH motif, with residues K404 and C405, and the hydrophobic loop (Φ-loop) residues N361, N363. These motifs are common in SF3 helicases and have been found to participate in both DNA melting and DNA unwinding [46–48]. A critical residue in the Φ-loop motif is a conserved phenylalanine residue that promotes DNA unwinding by intercalating and stacking with DNA bases [46]. Several other proteins use phenylalanine and other bulky residues to distort the DNA structure; the best examples include TBP, TFAM, and Sox 4 [49–51]. Studies have shown that hydrophobic residues are universal elements for DNA-structure deformation, particularly the widening of minor grooves [52]. It is not too difficult to visualize a possible Rep68-induced DNA melting mechanism where three consecutive F364 residues act as a hydrophobic wedge to initiate DNA melting. We showed that mutation F364A drastically affects the ability of Rep68 to unwind and nick DNA (Figure S5), but can DNA melting be induced by just three or a subset of Rep68 subunits? There is precedent based on studies of the origin of replication melting by the Papilloma Virus (PV) E1 helicase. PV DNA replication is contingent on assembling an E1 double hexameric complex (DH) on the origin of replication [53]. Studies have shown that the formation of the DH is a stepwise process dependent on DNA melting, with conclusive data showing that an E1 double trimer (DT) intermediate catalyzes the DNA melting process [54, 55]. The process is initiated by a local DNA distortion generated by the action of E1 PS1βH and Φ-loop residues and by the rotation of the two opposite trimers relative to each other to generate DNA untwisting [55]. In the AAV system, DNA melting could consist of the AAA^+^ domains undergoing continuous cycles of ATP hydrolysis, acting as crowbars inserted into the DNA to generate local melting (Figure 7). Although the DNA distortion we observe in our structure is small, the AAA^+^ domains can have additional movement, as observed in the Rep68-ssDNA hexameric complex structure [35]. The hexameric structure shows that the movement of the AAA^+^ domains may be large enough to potentially distort dsDNA to induce melting [35]. The limitation of our study is the use of ATPγS instead of ATP; however, this will require the capture of multiple snapshots along the ATP hydrolysis cycle, which we are currently working on. The study presented here is limited to Rep68 in a complex with an AAVS1 dsDNA substrate and does not address whether the structure of the complex will be similar in context to an ITR substrate. Nevertheless, taken together, our structures provide valuable views into how AAV Rep proteins are assembled on dsDNA and insights into DNA melting, further confirming the unique properties of this particular SF3 family member.

**Figure 7.**
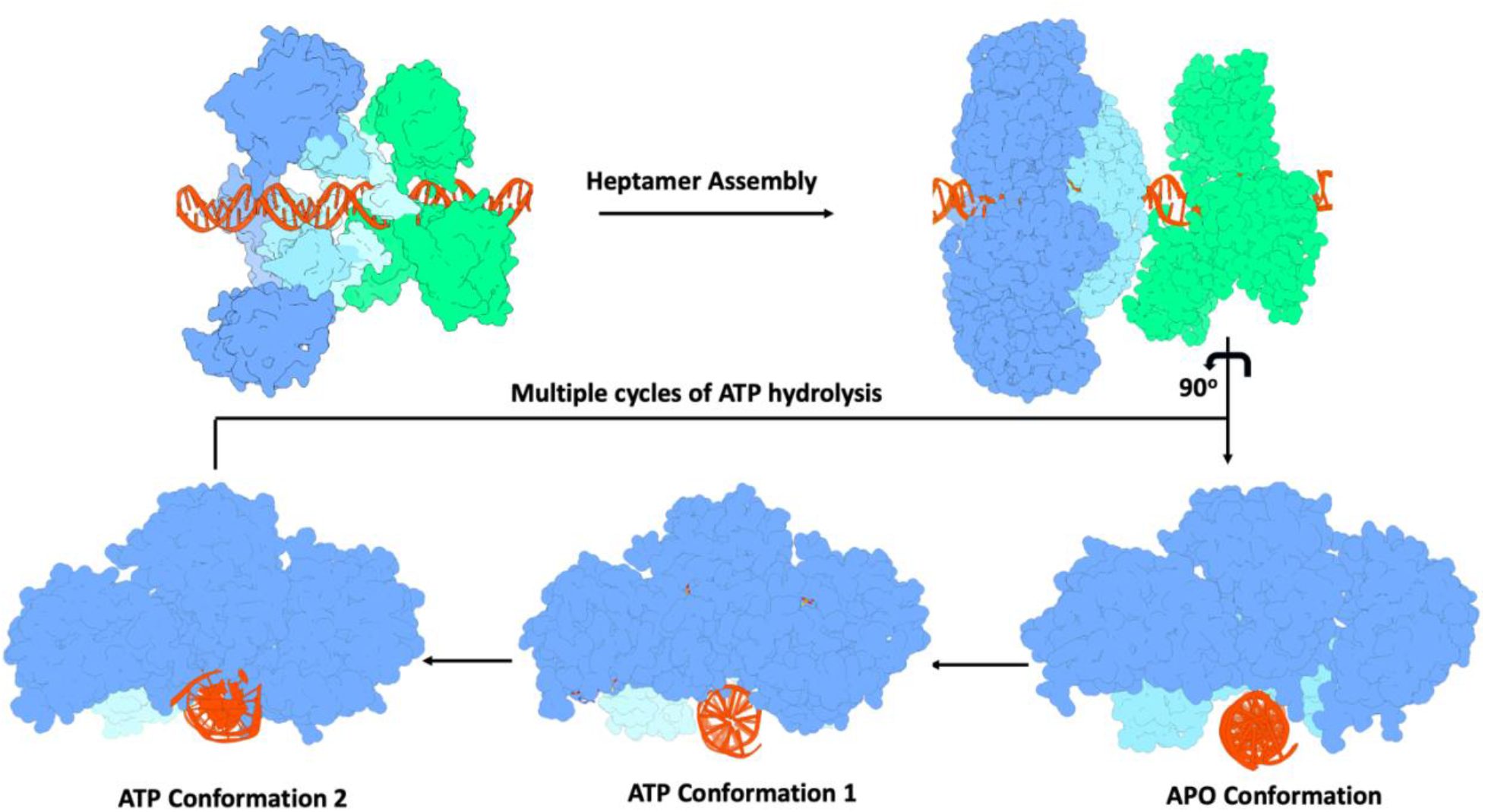
Proposed model of Rep68 assembly and DNA distortion. The OBDs interact with the RBS GCTC repeats to direct the initial assembly. The initial trimer further assembles into a heptamer that, upon multiple cycles of ATP binding and hydrolysis, could lead to DNA deformation and melting. Conformation 1 shows the structure of the Rep68-AAVS1 in this study. ATP conformation 2 shows that the AAA^+^ domains can further rotate, producing additional distortion.

## Material and Methods

### Cloning and Mutagenesis of Rep Expression Constructs

All mutant proteins were generated using the pHisRep68/15b plasmid containing the AAV2 Rep68 ORF (1-536) or the truncated form (1-490) were subcloned in vector PET-15b (Novagen). Site-directed mutagenesis for all mutants was generated using the QuickChange® mutagenesis kit (Stratagene). The sequences of all constructs were confirmed by DNA sequencing (Azenta).

## Protein expression and purification

All proteins were expressed using a modified pET-15b vector, expressed in E. coli BL21(DE3) cells (Novagen), and purified as described before [56]. Rep68*, a C151S mutant was used in all experiments and will be referred as Rep68 in the manuscript [32]. The final buffer contains (25 mM Tris-HCl [pH 8.0], 200 mM NaCl, and 2mM TCEP). His6-PreScission Protease (PP) was expressed in BL21(DE3)-pLysS at 37 °C for 3 h, in LB medium containing 1 mM IPTG. Cell pellets were lysed in Ni-Buffer A (20 mM Tris-HCl [pH 7.9 at 4 °C], 500 mM NaCl, 5 mM Imidazole, 10% glycerol, and 1 mM TCEP). After five 10-s cycles of sonication, the fusion protein was purified using a Ni-column – equilibrated in Ni-buffer A. Protein eluted was desalted using buffer A and a HiPrep^TM^ 26/10 desalting column (GE Healthcare). The hexahistidine tag was removed by PreScission protease treatment using 150 μg PP /mg His-PP-Rep68. During overnight incubation at 4 °C. Subsequent Ni-column chromatography using the buffer B (same as buffer A but with 1 M imidazole), was performed to remove the uncleaved fusion protein, and untagged Rep68 and was eluted with 50 mM imidazole. Rep68 was finally purified by gel filtration chromatography using a HiLoad Superdex 200 16/600 PG column (GE Healthcare) and Size Exclusion buffer (25 mM Tris, 200 mM NaCl, 1mM TCEP, pH 8.0). N-terminus His6-tagged WT and mutant Rep68 proteins were concentrated to 2 mg/ml with 50% glycerol, flash-frozen in liquid N_2_, and stored at -80°C.

### Supercoiled (sc) DNA nicking assay

ScDNA nicking activity for Rep68 was assayed as described previously [57]. Briefly, assays were performed in 30μl reactions containing 30mM Hepes-KOH (pH 7.5), 7mM MgCl2, 0.5mM DTT, 4mM ATP, 40mM creatine phosphate (Sigma), 1μg creatine phosphokinase (Sigma) in 15mM NaCl final concentration. 100ng scDNA plasmid and 200ng of purified Rep68 (or mutants) were added to the reactions. All samples were incubated for 1h at 37°C; the reaction was terminated by adding 10μl of stop reaction (proteinase K [1.2μg/μl], 0.5% SDS and 30mM EDTA pH7.5) and incubating for 1 h at 37°C. Samples were resolved in a 1% agarose gel (1X TAE), which was subsequently stained with ethidium bromide (0.3μg/ml) in 1X TAE. The scDNA plasmids used in this assay were pUC19 (containing an AAV2-ITR sequence with RBS and trs).

### Helicase Assay

The substrate used in this assay is a heteroduplex DNA consisting of an 18-bp duplex region with a 15-nucleotide 3’ tail at the bottom strand. The top strand has the sequence 5’-AGAGTACGGTAGGATATGAACCAGACACATGAT-3’; the bottom strand with sequence 5’-CATATCCTACCGTACTCT-F-3’ is labelled at the 3’ end with fluorescein and is released upon unwinding. We used the unlabeled bottom strand as a trap to prevent reannealing of the displaced fluorescent strand. All reactions were performed in a buffer containing 25 mM HEPES, 50mM NaCl (pH 7.0) in a total volume of 50μl. Rep68 at different concentrations was mixed with double stranded F-DNA at a final concentration of 1 μM and incubated for 15 min. Reaction was started by adding 5mM ATP-Mg and 2.5 μM trap DNA. Reaction was incubated at 25°C for 1 minute. EDTA was used to stop the reaction at a final concentration of 20μM. Aliquots of 10μl were loaded in a 12% bis-acrylamide gel (30%) (19:1) using 6X-loading dye (0.25 xylene cyanol FF, 30% glycerol). A Gel Doc EZ Imager was used to conduct densitometry and analysis of the bands. Background lane subtraction, white illumination and an activation time of 300 sec was used for the analysis.

### EM sample preparation

The Rep68-AAVS1 was mixed with a 50-mer AAVS1 dsDNA on a Superdex 200 10/300 gel filtration column (Cytiva Life Sciences) pre-equilibrated using a buffer containing 10 mM Na(PO_4_)_2_ pH 7.0, 150 mM NaCl 1 mM TCEP. The complex was then concentrated to 2 mg/ml using Amicon Ultra-4 centrifugal filters (Millipore). C-Flat carbon grids CF1.2/1.3-4C, 400 mesh Cu (Electron Microscopy Sciences CF413-50) were glow-discharged for 45 seconds with amylamine using a PELCO easiGlow™ glow-discharge system. For the ATPγS complex, 5mM MgCl_2_ and 5mM ATP_γ_S (final concentration) were added to the complex and incubated at room temperature for 5 minutes just before spotting the sample.

### Cryo-EM grid preparation and data collection

C-flat grids were glow-discharged for 40 seconds with amylamine using a PELCO easiGlow™ glow-discharge unit and spotted with 3.0 µL of a 0.5 mg/ml-2mg/ml protein complex. The sample was manually blotted for 1.5 seconds using a Vitrobot automatic cryo-plunge machine and plunged into liquid ethane. For storage, the samples were stored in liquid nitrogen. Initial screening was done on a Thermo Fisher Glacios TEM at the University of Virginia Molecular Electron Microscopy core facility and at the Pacific Northwest for Cryo-EM Center (PNCC).

#### (Apo complex)

Final data sets for the Apo complex were collected at the National Center for CryoEM Access and training (NCCAT) located at the New York Structural Biology Center on a Titan Krios operating at 300 kV at a nominal magnification of 81,000x with a Gatan K3 Summit direct detector with an imaging energy filter (Calibrated to a pixel size of 0.5291 Å). A total of 15602 image stacks were obtained with a defocus range of -0.6 to -2.9 μm. Each stack movie with 50 frames was recorded for a total of 2.5 s (0.05 s per frame) for a total dose of 65.58 electrons per Å^2^.

#### (ATPγS complex)

The data sets were collected at PNCC on a Titan Krios K3 operating at 300 kV at a nominal magnification of 81,000x with a Gatan K3 Summit direct detector (Calibrated to a pixel size of 0.526 Å). A total of 8301 image stacks were obtained with a defocus range of 0.6 to -2.6 μm. Each stack movie with 70 frames was recorded for a total of 3.3 s for a total dose of 46 electrons per Å^2^.

### Cryo-EM image processing and model building

#### Rep68-AAVS1 (Apo)

All data was processed using cryoSPARC v4.4 [58]. A total of 15,602 movie stacks were analyzed. Micrograph frames were aligned and superimposed using Patch Motion correction and Contrast transfer function (CTF) parameters were estimated with patch CTF Estimation [58]. Initial particle picking was done with a subset of micrographs using an automated blob picker followed by 2D classification to generate templates for template-based particle picking. Finally, ∼3.4 M particles were automatically picked using Topaz [59]. False particles and junk classes were removed through multiple rounds of 2D classification and the initial 3D models were generated using the *ab initio* algorithm in cryoSPARC. False-positive selections and contaminants were excluded through iterative rounds of heterogeneous classification. A final set of 453,032 particles was refined using non-uniform refinement, resulting in a map with an overall resolution of 6.1 Å. The final particle set was subjected to reference motion correction (polishing) and were used for a final non-uniform refinement job to produce the final map with an overall resolution of 5.4 Å using FSC 0.143 cutoff. The reconstructions were further improved by employing the deep learning algorithm of DeepEMhancer [34].

#### Rep68-AAVS1(ATPγS)

All Cryo-EM data processing was performed in Cryosparc 4.4 (Structura Biotechnology Inc). A total of 8301 movie stacks were subjected to Patch motion correction. The CTF parameters were estimated using Patch CTF. Data were selected using CTF fitted better than 8 Å, resulting in 8183 micrographs. a set of 294,324 particles were visually inspected through a series of passes of 2D classification and 2D selection to remove junk and low-quality particles, resulting in 78,701 good particles. After generating an initial ab initio model and 3D non-uniform refinement, the final map had an overall resolution of 5.3 Å using the FSC 0.143 cutoff.

All molecular models were built with Chimera and Coot and refined with Phenix and Isolde [60–62]. For the apo complex the crystal structures of Rep68 AAV-2 OBD-AAVS1 X-ray structure (PDB:4ZQ9) and AAV-2 Rep40 (PDB:1S9H) were used to build Rep68 subunits. Initially, 7 individual Rep40s and a 50 bp model of AAVS1 were manually docked into the cryo-EM map and oriented using Fit in Map in Chimera. The resulting model of the heptameric HD was further refined using rigid body refinement in PHENIX. For the ATPγS complex, only the Rep40 model was used during the initial docking refinement in Phenix; two rigid bodies were defined for each helicase subunit. The rigid body 1 contained the OD (residues 1-278) and rigid body 2 the AAA^+^ domain (residues 279-490). Model geometry was improved using ISOLDE. All superposition analysis of the models was done using the program LSQMAN [63].

## Funding

CRE is supported by National Institute of Health (NIH) 2R01GM124204. AW was funded by the Postbaccalaureate Research Program (VCU PREP).

A portion of this research was carried out at the NCCAT and the Simons Electron Microscopy Center located at the New York Structural Biology Center, supported by the NIH Commond Fund Transformative High Resolution Cryo-Electron Microscopy program (U24 GM1 129539) and by grants from the Simons Foundation (SF349247) and NY State.

A portion of this research was supported by NIH grant U24GM129547, performed at the PNCC at OHSU, and accessed through EMSL (grid.436923.9), a DOE Office of Science User Facility sponsored by the Office of Biological and Environmental Research.

## Acknowledgments.

We would like to thank Ed Eng, Elina Kopylov, and the outstanding staff from the National Center for Cryo-EM Access and Training (NCCAT). We would like to acknowledge Lauren Hales Beck, Nancy Meyer from the Pacific Northwest Cryo-EM Center (PNCC) supported by NIH grant U24GM129547.

## Author Contributions

R.J., V.S., and C.R.E. designed the research; V.S., R.J., B.B., A.W., and C.R.E. performed the research; and R.J., V.S., and C.R.E. wrote the paper, with all authors providing comments or revisions.

## Notes

### Competing Interest Statement

The authors have declared no competing interest.

