## Supplementary Figures for "Cryo-EM Structure of AAV2 Rep68 bound to integration site AAVS1: Insights into the mechanism of DNA melting"

| Table S1 – Cryo-EM data collection, refinement and validation statistics | | |
|---|---|---|
| **Data collection and processing** | | |
| | **Apo** | **ATPγS** |
| Instrument | FEI Titan Krios / Gatan K3 Summit (NCCAT) | FEI Tital Krios / Gatan K3 Summit (PNCC) |
| Magnification | 85,000 | 81,000 |
| Voltage (kV) | 300 | 300 |
| Energy filter slit width (eV) | 20 | 20 |
| Electron exposure ($e^-/Å^2$) | 65.58 | 50 |
| Defocus range (μm) | -0.8 to -2.5 | -1.53933 to -2.1033 |
| Pixel size (Å) | 0.5295 | 0.526 |
| Reconstruction pixel size (Å) | 1.059 | 1.052 |
| Symmetry imposed | C1 | C1 |
| Number of images | 15602 | 8301 |
| Number of frames per image | 50 | 70 |
| Data collection software | Leginon | SerialEM |
| Motion correction software | cryoSPARC v3.01 Patch Motion Correction | cryoSPARC v3.3.2 Patch Motion Correction |
| Initial particles | 3,466,434 | |
| Final particles | 320,617 | 379,193 |
| Map resolution at FSC threshold 0.146 (Å) | 5.3 | 3.4 |
| **Refinement** | | |
| **Model composition** | | |
| Non-hydrogen atoms | 52690 | 31657 |
| Protein residues | 3133 | 1943 |
| DNA | 82 | 42 |
| **ADP (B factors, min, max, mean ($Å^2$))** | | |
| Protein | 196.59/ 662.85/ 353.97 | 40.55/ 236.03/ 118.76 |
| DNA | 253.24/ 480.52/ 323.43 | 73.68/ 234.17/ 148.71 |
| Ligand | | 53.34/ 164.70/ 89.78 |
| **Bonds (RMSD)** | | |
| Bond lengths (Å) | 0.003 | 0.003 |
| Bond angles (°) | 0.833 | 0.559 |
| **Model-vs data** | | |
| CC (mask) | 0.62 | 0.7 |
| CC (box) | 0.73 | 0.66 |
| **Resolution estimates (Å)** | | |
| FSC model (0.143/0.5) Masked | 6.3/8.7 | 2.0/3.8 |
| FSC model (0.143/0.5) Unmasked | 6.3/2.8 | 2.0/3.8 |
| **Validation** | | |
| MolProbity score | 1.50 | 1.62 |
| Clashscore | 2.38 | 7.48 |
| Rotamer outlier (%) | 0.07 | 1.14 |

| Ramachandran plot (%) | | |
|---|---|---|
| Favored (%) | 92.08 | 97.10 |
| Allowed (%) | 7.53 | 2.90 |
| Disallowed (%) | 0.39 | 0.0 |

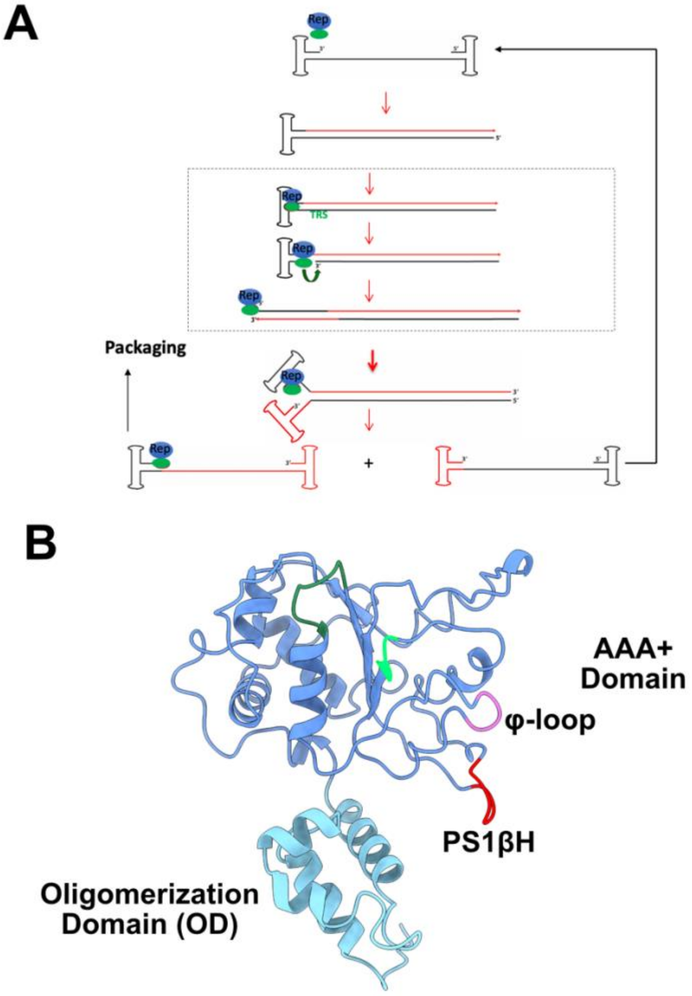


Figure S1. (A) Model of AAV DNA replication. The newly synthesized DNA is represented in red, with the 3'-end serving as a primer. The boxed region represents the terminal resolution reaction. Rep68 has the catalytic activities to bind, unwind, and nick at the terminal resolution site (trs) region. After nicking, Rep68 remains covalently bound to the 5'-end. The figure is a modification from Brister and Muzyczszka [18]. (B) Ribbon representation of AAV-2 Rep68 Helicase domain (HD)and functional motifs. Colored in sky blue is the Oligomerization domain (residues 215-278); colored in cornflower blue is the AAA$^{+}$ domain. Functional domains: sea green (Walker A, P-loop); spring green, Walker B; red, pre-sensor1 β-hairpin; hydrophobic loop (φ-loop), orchid.

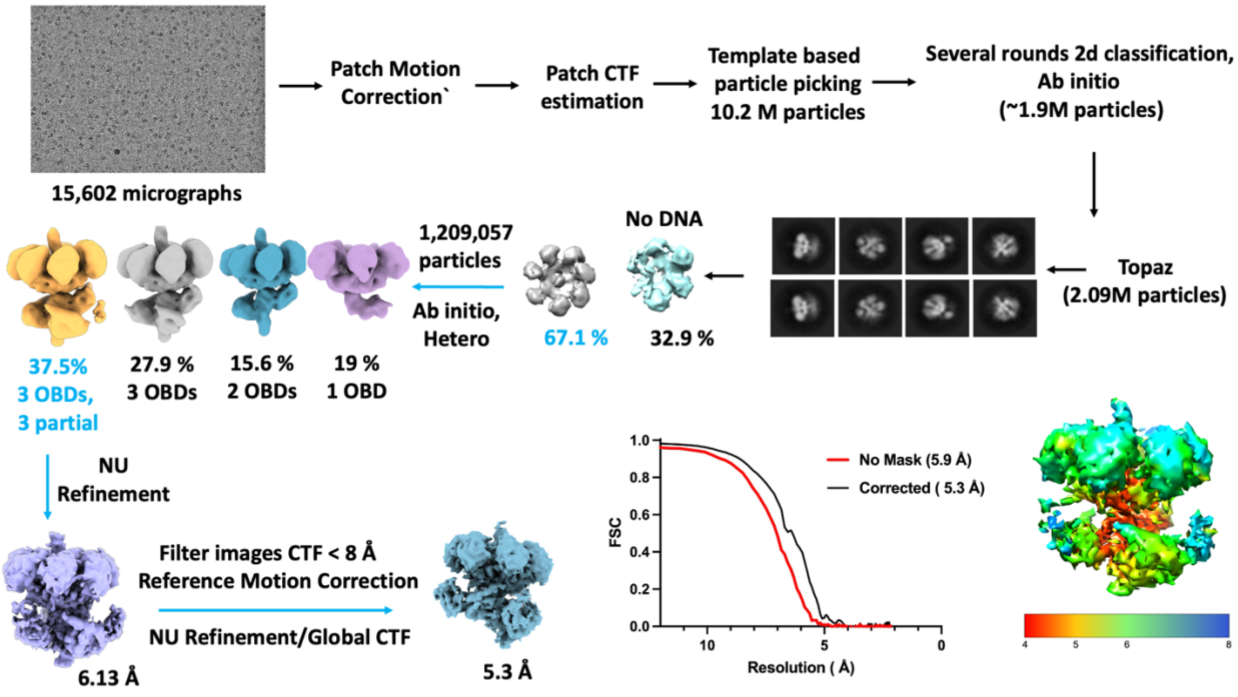


Figure S2. Cryo-EM data processing pipeline of the apo Rep68-AAVS1 complex.

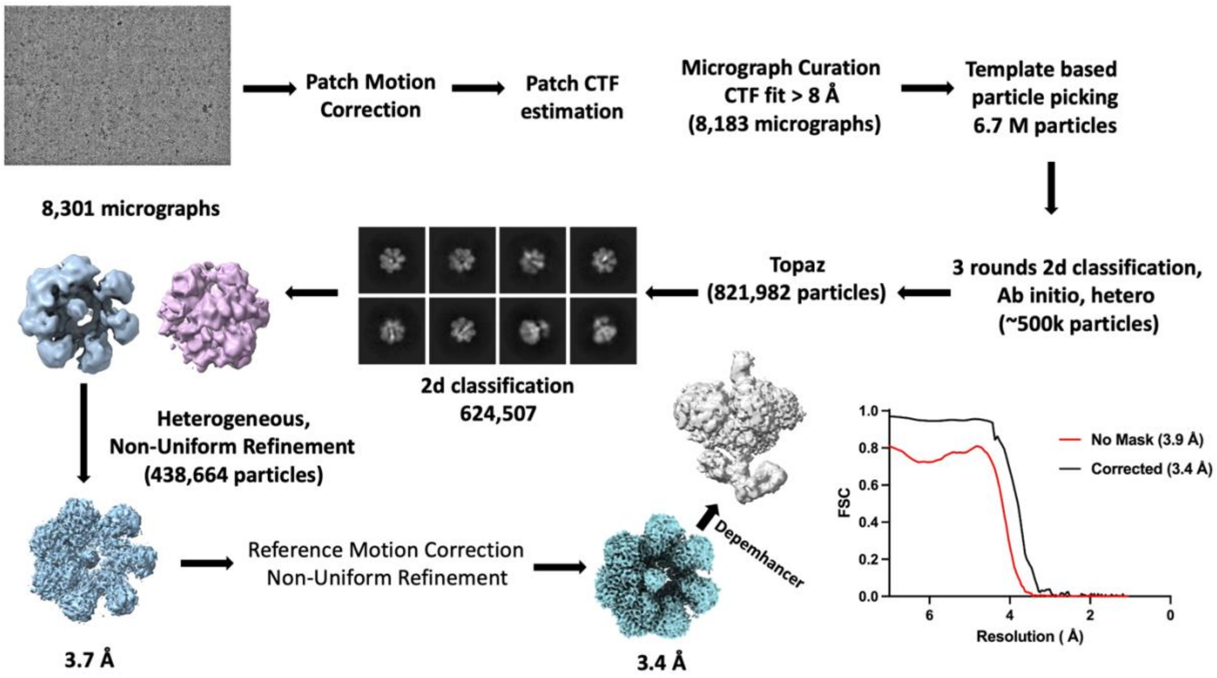


Figure S3. Cryo-EM data processing pipeline of the ATPγS Rep68-AAVS1 complex.

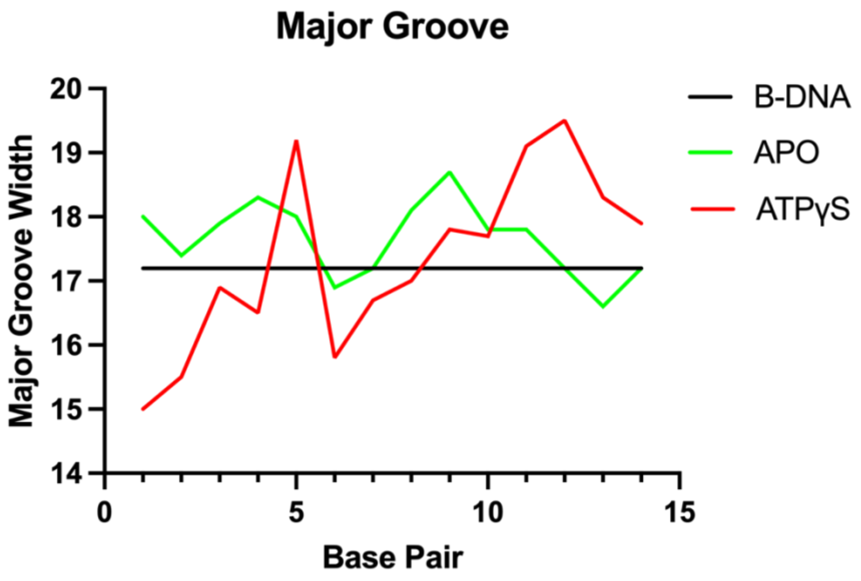


Figure S4. Comparison of DNA Major groove width between B-AAVS1 DNA (black), APO-AAVS1 (green) and ATPγS complex AAVS1 (red).
